# Changes in corticospinal excitability associated with motor learning by observing

**DOI:** 10.1101/205385

**Authors:** Heather R. McGregor, Michael Vesia, Cricia Rinchon, Robert Chen, Paul L. Gribble

## Abstract

While many of our motor skills are acquired through physical practice, we can also learn how to make movements by observing others. For example, individuals can learn how to reach in novel dynamical environments (‘force fields’, FF) by observing the movements of a tutor. Previous neurophysiology and neuroimaging studies in humans suggest a role for the motor system in motor learning by observing. Here we tested the role of primary motor cortex (M1) in motor learning by observing. We used single-pulse transcranial magnetic stimulation (TMS) to elicit motor evoked potentials (MEPs) in right hand muscles at rest. MEPs were elicited before and after participants observed either a video adapting her reaches to a FF or a control video showing a tutor performing reaches in an unlearnable FF. We predicted that observing motor learning would increase M1 excitability to a greater extent than observing movements that did not involve learning. We found that observing FF learning increased MEP amplitudes recorded from right first dorsal interosseous (FDI) and right abductor pollicis brevis (APB) muscles. There were no changes in MEP amplitudes for control participants who observed a tutor performing reaches in an unlearnable, randomly varying FF. The observed MEP changes can thus be specifically linked to observing motor learning. These results are consistent with the idea that observing motor learning produces functional changes in M1, or corticospinal networks or both.

## Introduction

Action observation activates brain areas involved in movement production. For example, in the macaque, so-called mirror neurons in area F5 of the premotor cortex are active both while the monkey performs goal-directed actions and while observing others performing similar actions (Di Pellegrino et al. 1992; Gallese et al. 1996; Rizzolatti et al. 1996). In humans, neuroimaging and neurophysiological studies have similarly shown that action observation engages the motor system (Strafella and Paus 2000a; Buccino et al. 2001; Watkins et al. 2003). The majority of this research has focused on the potential role of observation-related brain activity in higher cognitive functions such as action recognition, understanding others’ intentions, imitation, autism, theory of mind, empathy, etc. (Gallese 1998; Prinz 2006). However, a growing body of work has demonstrated that action observation can also facilitate motor learning.

Observation-related gains in motor performance have been reported using various experimental paradigms. For example, individuals can learn implicit button press sequences from observing others performing a serial reaction time task (Heyes and Foster 2002), learn novel dance sequences using observation-based training (Cross et al. 2006), and learn about object weights from observing others’ lifts (Alaerts et al. 2010; Buckingham et al. 2014). Of most relevance to the current study is the finding that participants can learn about how to reach in novel force environments from observing the movements of others (Mattar and Gribble 2005). In this study, participants observed a video of another individual (‘a tutor’) adapting his reaching movements to a force field (‘FF’) which was applied by a robotic arm. Participants who later performed reaches in the same FF as that which they had observed in the video showed a benefit, performing straighter movements in the FF compared to non-observed control participants. Conversely, participants who later performed reaches in the opposite FF to what they had observed showed a detriment, performing more curved movements in the FF compared to non-observing control participants. This study therefore demonstrated that participants were able to learn something about how to reach in novel FF environments through observing the movements of others. (Mattar and Gribble 2005) further found that performing an unrelated bilateral arm movement task during the video reduced the extent to which participants learned from observation, while performing a cognitive distractor task during observation did not. This finding suggested that motor learning by observing depends on the engagement of the observer’s motor system.

Recent neuroimaging and repetitive transcranial magnetic stimulation (rTMS) studies suggest a role for the motor system in motor learning by observing. Using resting-state fMRI, we have previously shown that observing FF learning changes functional connectivity between visual area V5/MT, the cerebellum, S1, and M1. Observation-related functional connectivity changes within this network were correlated with subsequent behavioral measures of motor learning by observing (McGregor and Gribble 2015). This study suggested that observing motor learning results in functional changes among visual and sensory-motor brain areas. Brown and colleagues ((Brown et al. 2009) investigated the role of M1 in motor learning by observing using low frequency repetitive transcranial magnetic stimulation (rTMS). In this study, participants observed a video of a tutor adapting to a FF and then received rTMS to left M1 following observation in order to reduce M1 excitability. Reducing M1 excitability following observation disrupted motor learning by observing. Participants who received rTMS to M1 and were later tested in the same FF to what was observed did not show behavioral gains associated with observation. Participants who received rTMS to M1 and were then tested in the opposite FF to what was observed did not show interference in the behavioral assessment. These results suggest that M1 plays a key role in motor learning by observing.

In the present study, we tested the role of primary motor cortex (M1) in motor learning by observing using TMS to probe for changes in corticospinal excitability following the observation of learning. Motor evoked potentials (MEPs) were elicited from muscles in the right hand before and after participants observed either a video depicting a tutor adapting her reaches to a FF or a control video showing a tutor performing reaches in an unlearnable FF. If M1 is indeed involved in motor learning by observing, then observing motor learning should increase M1 excitability to a greater extent than observing movements that did not involve learning. As predicted, we found that observing FF learning was accompanied by increases in MEP amplitudes. No changes in MEP amplitudes were seen for participants who observed a tutor performing reaches in an unlearnable FF. The MEP changes reported here are thus not due to action observation in general or observation of movement errors, but rather can be specifically linked to observation of motor learning. These results provide further support for the idea that observing motor learning involves functional changes in M1 or corticospinal networks, or both.

## Methods

### Participants

A total of 32 healthy volunteers participated in this study: 16 participants observed a video depicting a tutor learning to reach in a clockwise force field (5 males, mean age 21.6 year ± 0.71 SEM), and 16 participants observed a control video that depicted the same kinds of curved movements but did not depict learning (4 males, mean age 21.3 year ± 0.76 SEM). All participants were right handed, had normal or corrected-to-normal vision, and were naïve to force fields. None of the participants had neurological disorders, musculoskeletal disorders, or any contraindications to TMS (Keel, 2001). Participants provided written informed consent to experimental procedures approved by the Research Ethics Board at the University of Western Ontario.

### Reaching Task

Participants were seated in front of a custom tabletop and grasped the handle of a two degree-of-freedom robotic arm (IMT2, Interactive Motion Technologies) with the right hand (as shown in Figure 1A). The participant’s upper arm was abducted approximately 90^º^ from the trunk. The right arm was supported against gravity by an arm sled secured beneath the upper arm. An LCD TV projected visual feedback onto a semi-silvered mirror mounted horizontally above the robotic arm during the reaching task.

Participants were instructed to guide the robot handle from a central start position to a visual target, and were told to move the hand in a straight line. Eight targets were spaced equally around the circumference of a circle (see Figure 1A). Each target was a white circle (24 mm in diameter) located 10 cm from the start position (represented by blue circle that was 20 mm in diameter). A 5-mm pink circular cursor represented the position of the hand. To keep movement speed consistent from trial-to-trial, participants were provided with movement timing feedback at the end of each trial. The target turned blue if the movement duration was within the desired time range of 375 ± 100 ms, turned red if the movement was too fast, or turned green if the movement was too slow. Following movement timing feedback, the start position (blue circle) reappeared to signal that the participant should move the robot handle back to the start position to begin the next trial.

The robotic arm applied a velocity-dependent force field (FF) during the reaching task according to the following equation:

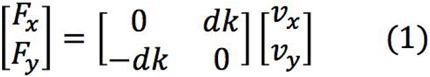

in which x and y are lateral and sagittal directions, Fx and Fy are the robot forces applied at the hand, v_x_ and v_y_ are hand velocities, k=15 Ns/m, and d=+1 (CW FF), −1 (CCW FF) or 0 (null field).

**Figure 1.**
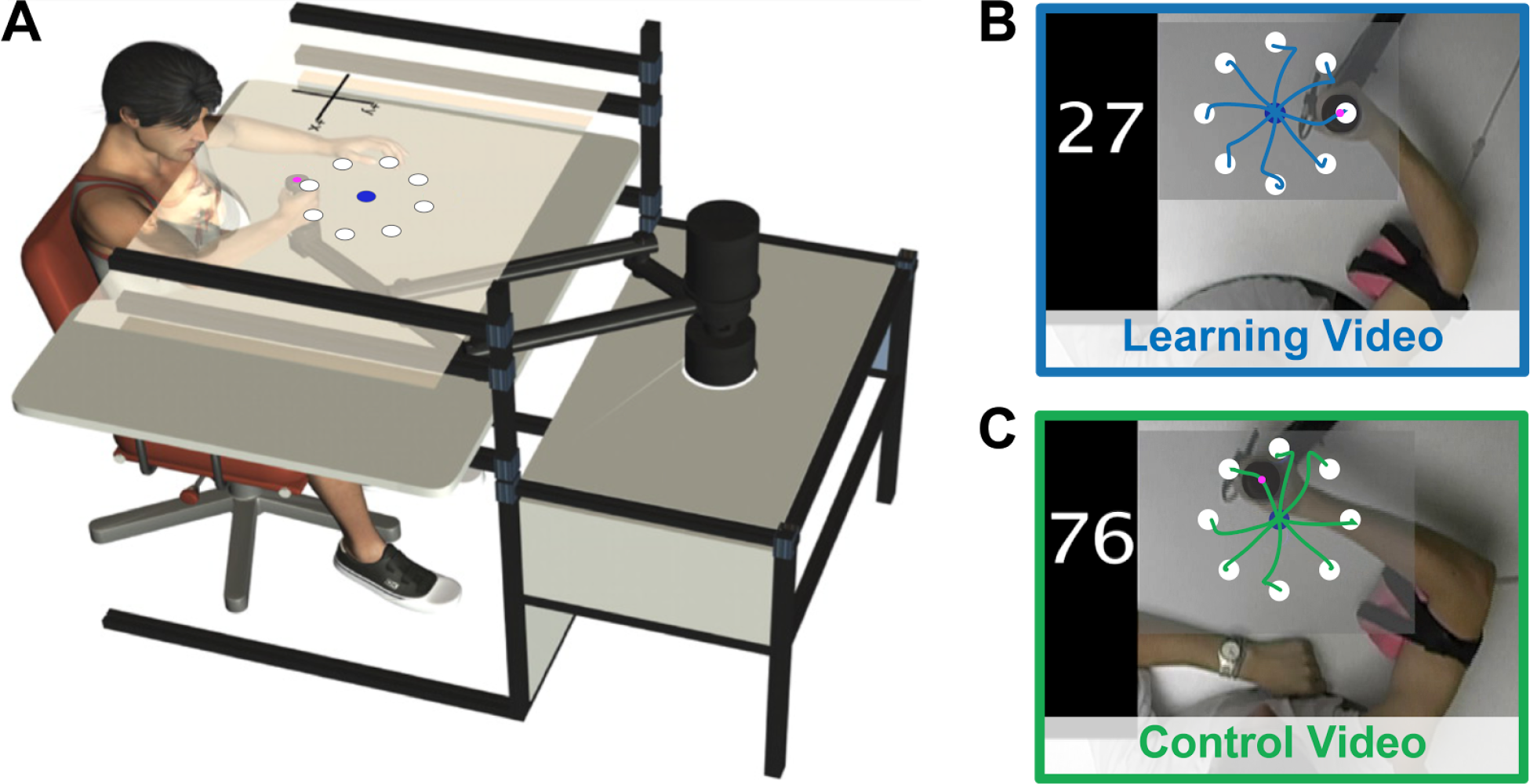
Task. **A) Reaching task.** Participants grasped the handle at the end of a robotic arm and performed straight reaches from a central start position (blue circle) to 8 targets (white circles). **B) Screenshot of the learning video.** The learning video showed a tutor adapting his reaches to a clockwise force field (CW FF). Superimposed trajectories are for demonstrative purposes only. **C) Screenshot of the control video.** The control video showed a tutor performing reaches in an unlearnable FF in which the direction of the applied force varied randomly from trial-to-trial. Superimposed trajectories are for demonstrative purposes only. FF, force field.

### Reaching Video Stimuli

Two videos were used in the study, and each one showed a tutor performing the reaching task described above from a top-down perspective. Both of the tutors were naïve to force fields. One video depicted a tutor adapting his reaches to a clockwise force field (CW FF) over 96 trials. The video depicted highly curved movements that gradually straightened as the tutor adapted to the CW FF. Participants observed this video twice (12 minutes total, 192 reaches observed in total). A screenshot of the video is shown in Figure 1B. Note that superimposed trajectories have been included for demonstrative purposes but were not shown to participants. Participants in a control group were shown a video that depicted a tutor performing 96 reaches in an unlearnable FF. In the unlearnable FF, the direction of the force field varied randomly from trial-to-trial between a CCW FF, CW FF or null field. The control video therefore showed the tutor performing both high and low curvature movements, but lacked the progressive decrease in movement curvature that was depicted in the learning video. Participants assigned to the control group observed the control video twice (12 minutes total, 192 reaches observed in total). A screenshot of the control video is shown in Figure 1C. Note that superimposed trajectories were not shown to participants.

### Experimental Design

The experimental design is shown in Figure 2A. Participants were first familiarized with the robotic arm and the reaching task by performing approximately 40 practice reaches in a null field. Then, in a baseline condition, participants performed 96 reaches in a null field (12 reaches to each of the 8 targets). This was done to assess the participant’s baseline movement curvature prior to observation.

Corticospinal excitability was assessed before and after observation using single-pulse TMS (see below for details). We recorded motor evoked potentials (MEPs) from muscles in the right hand both before and after observation of either the learning video or the control video. We chose to examine MEP changes involving hand muscles for two reasons: first, we have found in previous work that observing motor learning changes functional connectivity between the the hand area of M1, primary somatosensory cortex, the cerebellum and visual area V5/MT (McGregor and Gribble 2015). Second, MEPs are more easily evoked from distal muscles compared to more proximal muscles (Palmer and Ashby 1992).Fifteen pre-observation MEPs were acquired from the relaxed first dorsal interosseus (FDI) and abductor pollicis brevis (APB) muscles in the right hand (see below for details). During MEP acquisition, participants were instructed to fixate a crosshair presented in their line of view while resting their forearms on a tabletop.

Next, one group of participants (n=16) observed the learning video showing a tutor adapting to a CW FF. A control group (n=16) observed the control video showing a tutor performing reaches in an unlearnable FF. In both cases, participants were not told about FFs in the videos. Participants were asked to count the number of times the tutor in the video performed a correctly-timed reach indicated by the target turning blue following a reach. This was done to verify that the participant paid attention to the videos. Following the videos, 15 post-observation MEPs were acquired from the right FDI and APB using the same stimulation site and intensity as in the pre-observation MEP acquisition.

As a behavioral test of motor learning by observing, all participants then performed 96 reaches in a CCW FF. Note that participants were tested in a FF opposite to the one that was observed in the learning video. The idea is that the more participants learned about the CW FF during observation, the more interference this would cause in the CCW FF. Therefore, greater motor learning by observing was indicated by greater movement curvature in the test CCW FF. As we have frequently done in past work (Mattar and Gribble 2005; Cothros et al. 2006; Brown et al. 2009; McGregor and Gribble 2015, 2017; McGregor et al. 2016), we chose to use an interference paradigm because this tends to yield a more sensitive measure of motor learning by observing.

**Figure 2.**
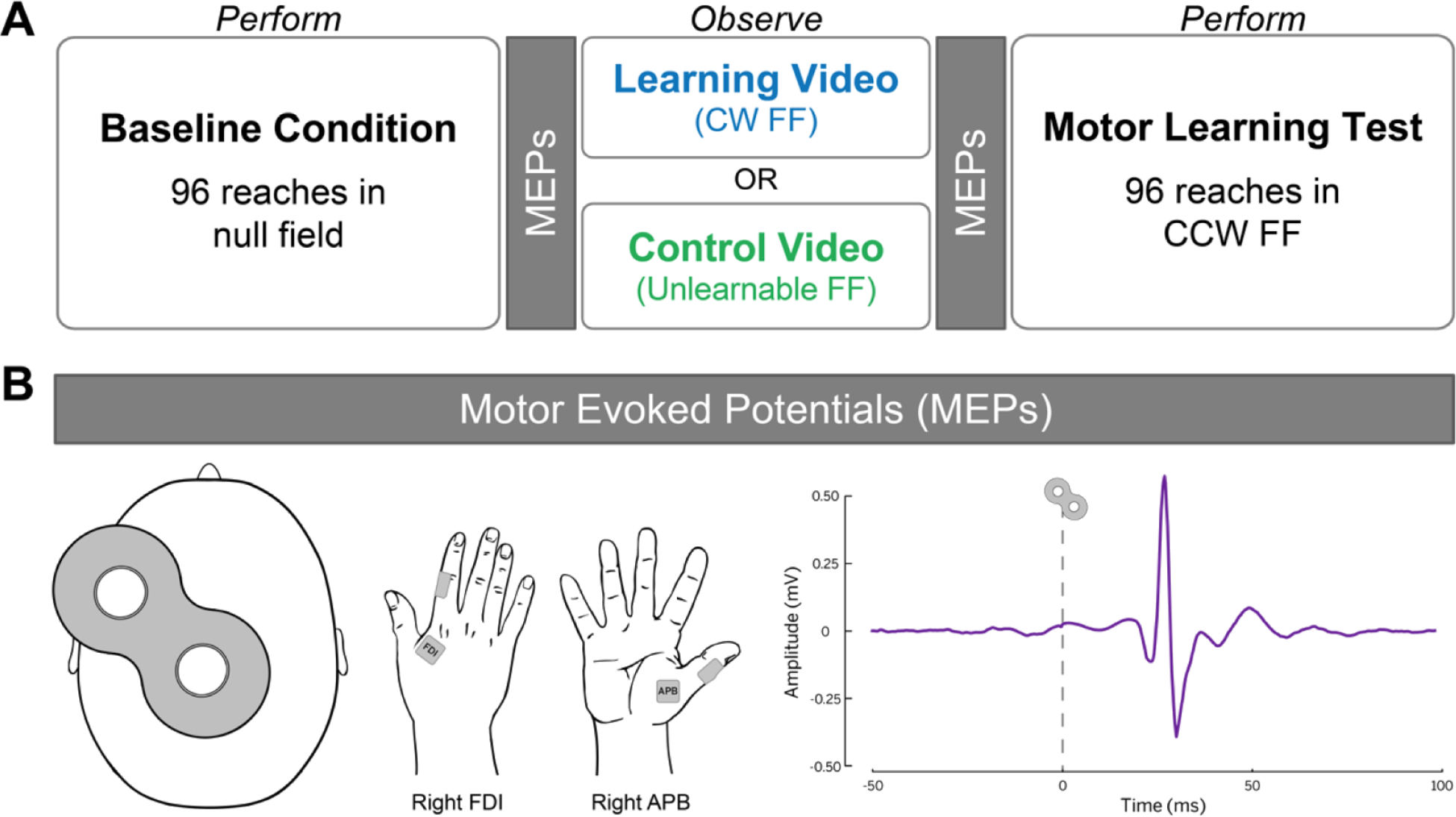
**A) Experimental design.** In the baseline condition, participants performed reaches in a null field in which the robotic arm applied no force. Fifteen pre-observation MEPs were acquired from the right FDI and APB muscles at rest. Participants then observed either a learning video or a control video. The learning video showed a tutor adapting his reaches to a clockwise force field (CW FF). The control video showed a tutor performing reaches in an unlearnable FF in which the direction of the applied force varied randomly from trial-to-trial. Fifteen post-observation MEPs were then acquired from the right FDI and APB muscles at rest. In the motor learning test, all participants performed reaches in a counterclockwise force field (CCW FF). **B) MEPs.** Motor evoked potentials (MEPs) were elicited by applying single-pulse TMS to the hand muscle representation of left M1 (shown at far left). MEPs were recorded from the relaxed first dorsal interosseous (FDI) and the abductor pollicis brevis (APB) muscles in the right hand (shown in middle). A sample MEP from one participant is shown on the far right. MEPs, motor evoked potentials; FDI, first dorsal interosseous; APB, abductor pollicis brevis; FF, force field; M1, primary motor cortex.

### Motor Evoked Potential (MEP) Acquisition

We probed corticospinal excitability by using single-pulse TMS to elicit motor evoked potentials (MEPs) before and after observation. Single monophasic TMS pulses were delivered using a custom 50-mm diameter figure-of-eight branding iron coil connected to a Magstim 200 mono pulse stimulator (Magstim, Whitland, UK). The TMS coil was placed on the scalp over the hand representation of left M1 and positioned 45° relative to the sagittal plane to induce a current in the posterior-to-anterior direction. This coil orientation likely activates corticospinal neurons trans-synaptically (Di Lazzaro et al. 2008). Single TMS pulses were delivered at a frequency of 0.25 Hz. MEPs were recorded using Ag-AgCl surface electrodes placed in a belly-tendon montage over the first dorsal interosseous (FDI) and the abductor pollicis brevis (APB) muscles in the right hand (see Figure 2B). Electromyographic (EMG) signals were amplified (1000x), bandpass filtered online (2 Hz - 2.5 kHz; Intronix Technologies Model 2024F, Bolton, Ontario, Canada), and digitized at 5 kHz by an analog-to-digital interface (Micro 1401, Cambridge Electronic Design, Cambridge, UK), and stored on a computer for offline analysis.

Prior to MEP acquisition, we identified each participant’s motor hotspot. The motor hotspot was defined as the location at which we could elicit MEPs of at least 50 mV in (peak-to-peak) amplitude from both the FDI and APB muscles in the relaxed right hand in at least 5 of 10 trials with the lowest stimulator intensity. After identifying the hotspot, we adjusted the stimulation intensity such that TMS pulses would elicit MEPs of approximately 1 mV (peak-to-peak amplitude) from the relaxed FDI and APB muscles. The same hotspot and stimulation intensity were used before and after observation within a given experimental session. We used a frameless stereotaxic neuronavigation system (Brainsight, Rogue Research, Montreal, QC, Canada) to ensure consistency in the TMS coil position.

### Behavioral Analysis

Behavioral data were collected at a sampling rate of 600 Hz. Data were lowpass filtered offline at 40 Hz. Each movement trajectory was rotated to a common axis such that it was aligned with the 0° target (straight ahead of the home position). We quantified the curvature of each movement by computing the maximum perpendicular deviation (PD) of the hand path. The PD of a movement was defined as the maximum point of lateral deviation of the hand path relative to a straight line connecting the home and (0°) target. Since movements were aligned to a common axis (aligned to the 0° target), reaches in the CCW FF were curved to the left, reflected as negative PD values. We then computed a measure of initial PD in the CCW FF for each participant. Initial PD in the CCW FF was computed as the average PD of the participant’s first 24 reaches (first 3 reaches to each of the 8 targets) minus the average PD of the last 48 reaches in the null field. This allowed us to assess the extent to which observation interfered with the participant’s initial performance in the CCW FF relative to his or her baseline movement curvature in the null field. As we have demonstrated previously (Mattar and Gribble 2005; Cothros et al. 2006; Brown et al. 2009; McGregor and Gribble 2015, 2017; McGregor et al. 2016), learning about a FF from observation results in more curved movements in the opposite FF. Therefore, we predicted that greater motor learning by observing would result in greater movement curvature in the CCW FF, and hence larger negative PD values.

### MEP Data Analysis

EMG data were bandpass filtered offline between 20-500 Hz and a notch filter was applied (58-62 Hz). The dependent measure in statistical tests was the peak-to-peak amplitude of MEPs. A 2×2×2 split-plot analysis of variance (ANOVA) was carried out using group (learning, control) and muscle (FDI, APB) as between-subjects factors, and time relative to observation (pre-observation, post-observation) as a within-subject factor.

## Results

### Behavioral Results

Figure 3A shows the evolution of PD over the course of the experiment for the learning and control groups. It can be seen that movements are straight in the baseline null field condition for both groups. Following observation, the learning group’s initial movements in the CCW FF are more curved compared to those of the control group. This difference in initial performance in the CCW FF can also be seen in Figure 3B in which we have shown hand path traces of movements performed in the first block of trials from individual participants, one from the learning group and the other from the control group. Therefore, as predicted, participants who observed the video of the tutor learning to reach in the CW FF experienced interference during initial performance in the CCW FF compared to control participants who did not observe learning. The two groups’ learning curves converge after the first few blocks of movements in the CCW FF. This is expected as participants in both groups begin to actively adapt to the CCW FF. Figure 3C shows initial PD in the CCW FF for both groups. Participants who observed the learning video (blue) exhibited greater movement curvature in the CCW FF and this was reflected as more negative PD values compared to participants who observed the control video (green) [t(30)=1.85, p = 0.037, one-tailed]. Therefore, as a result of having observed CW FF learning in the video, participants in the learning group performed worse, more curved movements in the CCW FF relative to the control group.

**Figure 3.**
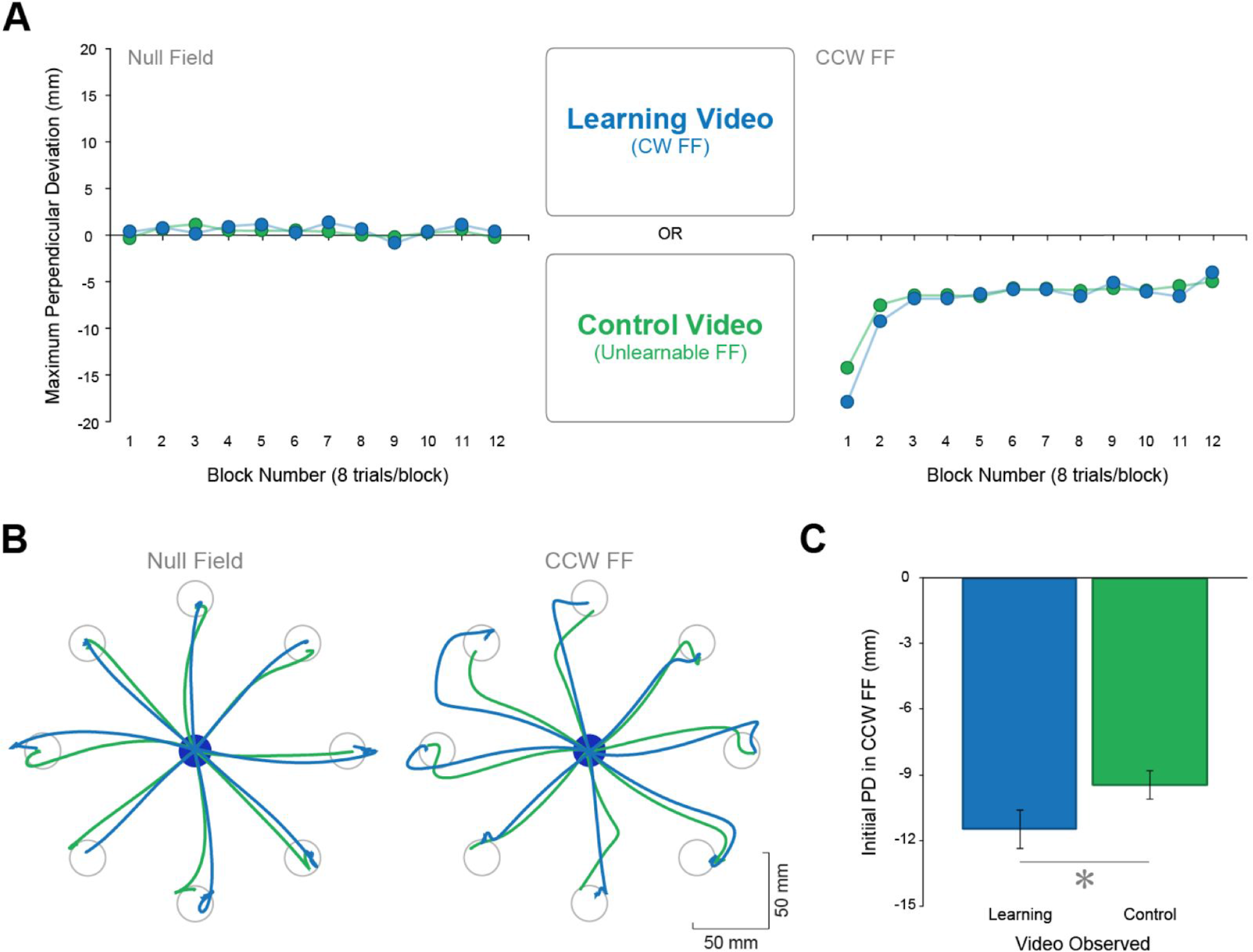
Behavioral Results. **A) Reaching behavior.** Evolution of movement curvature over the course of the experiment. Positive values along the y-axis indicate rightward curvature, negative values along the y-axis indicate leftward curvature. **B)** Sample trajectories. Typical hand trajectories in the null field (left) and in the CCW FF (right) from a participant in the learning group (blue) and a participant in the control group (green). **C)** Initial PD in the CCW FF. Average PD during the first 3 blocks in the CCW FF relative to the participant’s own baseline PD in the null field for the learning group (blue) and control group (green). * indicates p < 0.05. Error bars indicate standard error.

### MEP Results

We probed corticospinal excitability by using single-pulse TMS to elicit MEPs from hand muscles before and after participants observed either the learning video or the control video. A split-plot ANOVA revealed a statistically significant interaction between group (learning vs control) and time relative to observation (*F*(1,30) = 8.74, p < 0.01). Post-observation MEP amplitudes measured from both the FDI muscle and APB muscle increased for the group who observed the learning video (t(30) = 2.03, p < 0.05 and t(30) = 2.28, p < 0.05, Bonferroni-Holm corrected, respectively). This interaction is shown in Figure 4A which depicts post-observation MEP amplitudes as a percentage of pre-observation MEP amplitudes. It can be seen that greater increases in MEP amplitudes from pre-observation to post-observation were measured for the learning group compared to the control group, regardless of the hand muscle from which MEPs were recorded. This interaction effect is shown in greater detail in Figure 4B which shows pre-observation and post-observation MEP amplitudes for both groups and both hand muscles. Again, it can be seen that MEP amplitudes increased for both hand muscles following the observation of the learning video, but not following the observation of the control video.

**Figure 4.**
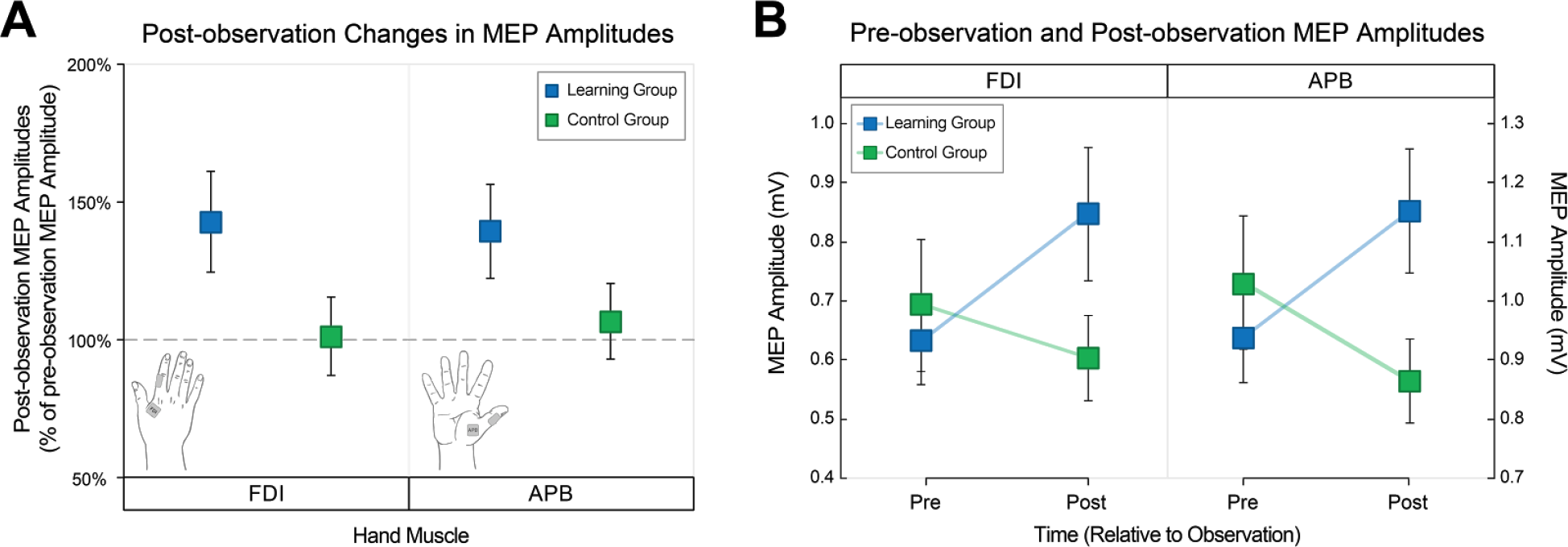
MEP Results. **A) Post-observation changes in MEP amplitudes.** Post-observation changes in MEP amplitudes reflected as a percentage of pre-observation MEP amplitudes. The horizontal, grey dashed line at 100% indicates no change in MEP amplitude from pre-observation. Data from the learning group and the control group are shown in blue and green, respectively. Post-observation MEP changes recorded from the FDI muscle are shown in the left panel. Post-observation MEP changes recorded from the APB muscle are shown in the right panel. **B) Pre- and post-observation MEP amplitudes.** Pre- and post-observation MEP amplitudes collected from FDI (left panel) and from APB (right panel). Data from the learning group and the control group are shown in blue and green, respectively. Error bars indicate standard error.

## Discussion

Here we used single-pulse TMS to test the role of M1 in motor learning by observing. We measured changes in corticospinal excitability by eliciting MEPs from the observer’s hand muscles before and after the observation of a FF reaching task. We found that those participants who observed the video of a tutor undergoing FF adaptation showed reliable increases in MEP amplitudes recorded from FDI and APB hand muscles. However, a control group who observed a video showing a tutor performing reaches in an unlearnable FF did not show post-observation changes in MEP amplitudes. These MEP changes can therefore be linked to observation of motor learning, and not to observing movements in general or observing movement errors. This result suggests that observation of motor learning involves functional changes in M1 or corticospinal networks or both.

Previous studies have examined how observing motor actions changes Ml excitability. However, here we examined how the observation of *motor learning* changes M1 excitability. Our finding that observing motor learning increases MEP amplitudes is consistent with other neurophysiological demonstrations of the facilitatory effect of action observation on neuroplasticity in M1 (Fadiga et al. 1995; Watkins et al. 2003; Stefan et al. 2005). For example, Stefan and colleagues (2005) have shown that observing the performance of simple thumb movements can alter motor memories encoded in Ml. Participants observed a video showing a tutor performing thumb flexion movements and a video showing thumb extension movements. During video breaks, single-pulse TMS was applied over the observer’s Ml and thumb twitches were elicited. It was found that TMS-evoked thumb movements were biased in the direction of the observed movements. For example, when a participant had observed a video showing thumb flexion, there was an increased probability that TMS-elicited thumbs movements would be in the flexion direction. When that same participant then observed a video showing thumb extensions, TMS-evoked thumb movements changed such that they were biased in the extension direction. These findings showed that motor memories encoded in primary motor cortex are subject to modification via action observation.

We found that observation of a tutor adapting to a consistent FF increased MEP amplitudes whereas observing a tutor performing reaches in a randomly-varying (and hence unpredictable) FF did not bring about MEP changes. A recent study performed by de Beukelaar and colleagues (2016) showed that being able to predict an upcoming observed movement can elicit anticipatory Ml facilitation. In this study, participants observed videos of an actor lifting an object using either a precision grip or a whole hand grip. Participants were provided with a cue prior to each video, informing them of the type of grip they would see the actor use in the upcoming trial. The video began and the actor’s arm hovered above the object, giving no visual indication of the grip type that would be used. During this pre-movement phase of the video, participants showed increases in MEP amplitudes from muscles that would be used in the predicted grasp. Therefore, being able to predict the upcoming observed movement resulted in anticipatory, muscle-specific Ml facilitation before the actor executed the movement (de Beukelaar et al. 2016). In the context of the current study, it is possible that this anticipatory facilitation may explain why we found MEP increases for the learning group but not for the control group. Recall that the upcoming force direction could only be predicted in the learning video, which depicted a CW FF applied throughout. It is feasible that participants who observed the learning video became able to predict the tutor’s upcoming movement kinematics and therefore showed a similar anticipatory Ml facilitation throughout the video. This may be a possible explanation as to why MEP amplitudes increased following the learning video only.

Here we found that observing FF learning, a reaching task primarily involving shoulder and elbow movement, facilitates MEPs acquired from hand muscles. This finding is at odds with previous work showing that changes in Ml excitability are specific to the muscles used for the observed movement (Fadiga et al. 1995; Strafella and Paus 2000b; Watkins et al. 2003; Stefan et al. 2005; Borroni and Baldissera 2008; Alaerts et al. 2010; de Beukelaar et al. 2016). For example, Strafella and Paus (2000) applied single-pulse TMS to M1 while participants observed videos showing an actor performing hand movements (i.e., writing) or arm movements (i.e., flexion, extension, abduction and drawing shapes with the arm while the wrist and hand remained a prone position). MEPs elicited from the observers’ hand muscles (FDI) increased during hand movement observation. MEPs elicited from the observers’ biceps increased during arm movement observation. Results from this study and others have shown that action observation facilitates the observer’s motor system in a muscle-specific manner.

We observed changes in MEP amplitudes from hand muscles following the observation of a learning task that largely involves shoulder and elbow movement. One possible reason why we observed changes in MEPs from hand muscles is that participants looked at the tutor’s hand and the robot handle during the video. While the reaching task involved elbow and shoulder movement, the goal of the task was to move the robot handle (and hand) to visual targets along a straight hand path. Looking at the tutor’s hand likely activates the hand area of the observer’s motor cortex, as we have indeed reported previously using similar video stimuli (McGregor and Gribble 2015). Moreover, while the FF reaching task used in the current study primarily involves changes in forces generated at the elbow and shoulder, there is evidence that grip forces also change with motor learning. For example, Flanagan and colleagues (2003) have shown that grip forces show anticipatory adjustments over the course of motor learning. In their experiment, participants gripped an object and were instructed to move it along a straight path. The object was attached to a robotic manipulandum which applied a FF that varied in magnitude with the velocity of movement. Participants learned to scale their grip forces well before they were able to accurately move the object along the desired path. Despite their use of a different grip type (precision vs. power grip) and FF (vertical versus horizontal curl field), this finding by Flanagan and colleagues (2003) may provide insight into why MEP changes were seen in hand muscles during observation of FF learning.

Here we showed that observing FF learning increases MEP amplitudes recorded from FDI and APB hand muscles. In contrast, control participants who observed a tutor performing reaches in an unlearnable, randomly-varying FF did not show post-observation changes in MEP amplitudes. This suggests that motor learning by observing involves functional changes in Ml or corticospinal networks or both. It is likely that projections from various brain regions such as premotor cortices mediate Ml activity during action observation and influence the extent to which visual input facilitates motor learning. This study contributes to growing evidence that motor learning by observing is supported by a broad network of visual, somatosensory and motor brains areas such as premotor cortex, primary motor cortex, primary cortex, superior parietal lobule, visual area V5/MT, and the cerebellum (McGregor and Gribble 20l5, 20l7).

## Acknowledgements

The authors wish to thank Dinant Kistemaker for the figure showing the robotic manipulandum setup. This work was supported by the Natural Sciences and Engineering Research Council of Canada, and by the National Institute of Child Health and Human Development R01 HD075740.

